# BIBSNet: A Deep Learning Baby Image Brain Segmentation Network for MRI Scans

**DOI:** 10.1101/2023.03.22.533696

**Authors:** Timothy J. Hendrickson, Paul Reiners, Lucille A. Moore, Jacob T. Lundquist, Begim Fayzullobekova, Anders J. Perrone, Erik G. Lee, Julia Moser, Trevor K.M Day, Dimitrios Alexopoulos, Martin Styner, Omid Kardan, Taylor A. Chamberlain, Anurima Mummaneni, Henrique A. Caldas, Brad Bower, Sally Stoyell, Tabitha Martin, Sooyeon Sung, Ermias A. Fair, Kenevan Carter, Jonathan Uriarte-Lopez, Amanda R. Rueter, Essa Yacoub, Monica D. Rosenberg, Christopher D. Smyser, Jed T. Elison, Alice Graham, Damien A. Fair, Eric Feczko

## Abstract

**Objectives:** Brain segmentation of infant magnetic resonance (MR) images is vitally important for studying typical and atypical brain development. The infant brain undergoes many changes throughout the first years of postnatal life, making tissue segmentation difficult for most existing algorithms. Here we introduce a deep neural network BIBSNet (**B**aby and **I**nfant **B**rain **S**egmentation Neural **Net**work), an open-source, community-driven model for robust and generalizable brain segmentation leveraging data augmentation and a large sample size of manually annotated images.

**Experimental Design:** Included in model training and testing were MR brain images from 90 participants with an age range of 0-8 months (median age 4.6 months). Using the BOBs repository of manually annotated real images along with synthetic segmentation images produced using SynthSeg, the model was trained using a 10-fold procedure. Model performance of segmentations was assessed by comparing BIBSNet, joint label fusion (JLF) inferred segmentation to ground truth segmentations using Dice Similarity Coefficient (DSC). Additionally, MR data along with the FreeSurfer compatible segmentations were processed with the DCAN labs infant-ABCD-BIDS processing pipeline from ground truth, JLF, and BIBSNet to further assess model performance on derivative data, including cortical thickness, resting state connectivity and brain region volumes.

**Principal Observations:** BIBSNet segmentations outperforms JLF across all regions based on DSC comparisons. Additionally, with processed derived metrics, BIBSNet segmentations outperforms JLF segmentations across nearly all metrics.

**Conclusions:** BIBSNet segmentation shows marked improvement over JLF across all age groups analyzed. The BIBSNet model is 600x faster compared to JLF, produces FreeSurfer-compatible segmentation labels, and can be easily included in other processing pipelines. BIBSNet provides a viable alternative for segmenting the brain in the earliest stages of development.

## Introduction

The first years of postnatal life are marked by significant neuronal development through synapse growth, myelination of axons, and programmed cell death ^1,2^. Converging evidence suggests that neurodevelopmental and psychiatric disorders throughout the lifespan are influenced by atypical brain development during this highly dynamic period ^3–7^. These dynamics and have led to major investments in characterizing brain trajectories during this sensitive period, including the recently launched Healthy Brain and Child Development (HBCD) Study™ - a large (N ∼ 7,200) multi-site, longitudinal, effort to examine the developing brain from 0-10y to study typical and atypical brain development.

Magnetic resonance imaging (MRI) is a non-invasive tool that can be used to study developmental brain health and disease. It is capable of producing multiple types of MR data, including diffusion, spectroscopy, functional, and quantitative MRI data ^8–11^. The vast majority of analysis strategies across all of these modalities depend on well-annotated, or segmented, MRI data - that is, data that delineates various brain tissues. Traditionally, structural MR images (i.e., T_1_-weighted (T1w) or T_2_-weighted (T2w) anatomical images) are used to create annotations that segment tissue types, such as white matter, gray matter, cerebrospinal fluid (CSF), and subcortical structures^12^. Accurate segmentation of cortical gray matter is also necessary to produce more advanced morphological metrics, such as cortical thickness, surface area, and gyrification^13^. Finally, other modalities such as functional MRI (fMRI) and diffusion MRI (dMRI) rely on accurate segmentations to produce more computationally sophisticated metrics like functional or structural connectivity ^14–18^. Studies such as HBCD require efficient and reliable methods to conduct this critical step required for many derived measures of MRI.

Automated brain segmentation algorithms often rely on high-resolution (e.g. ∼1mm^3^ or smaller) T1w and T2w anatomical images to annotate tissue types. These algorithms depend on voxel contrast and intensity differences across differing brain tissue and regions to delineate brain tissue and region boundaries. Brain tissue and region boundaries are, for the most part, easily delineated within the adult and child brain; however, they are often less accurate in infant data. This difficulty is due to the significant changes that the brain undergoes during the first years of postnatal life, such as myelination^1,19,20^. For example, T1w-gray matter voxel contrast is greater than T1w-white matter voxel contrast in infants 0 to 3-months (**Figure 3, Supplemental Figure 1**), T1w-gray matter and T1w-white matter are approximately equal in contrast from about 3-6 months causing the tissues to look very similar (**Figure 3, Supplemental Figure 1**), and at >6 months and on the T1w-gray matter is less than T1w-white matter, emulating the tissue contrast of an adult brain ^12,21,22^.

One popular approach in the field to produce accurate segmentations is joint label fusion (JLF) ^23,24^. JLF relies on manually annotated segmentations in multiple individual atlases. From there an individual’s brain is non-linearly registered to each atlas. A ‘winner take all’ approach is then used to assign labels to each voxel based on local cross correlations between voxel intensities in the subject and atlas. While this strategy has shown to be successful compared to single atlas approaches ^25,26^, the approach is time-consuming (computation of 2-3 days) and still error-prone. Furthermore, results are quite variable at different ages, which by themselves may take months to optimize. Exploring alternative solutions is warranted.

Convolutional Neural Networks (CNN) are an attractive alternative to traditional methods of segmentations, and have shown significant promise in adult samples^27^, CNNs are fast and take only minutes to segment an infant’s brain once properly trained. CNNs can be further trained with new data from other datasets in order to boost generalizability quickly. Prior work in infants has attempted to build CNNs for fast brain segmentation ^22,28–31,32^. **Supplemental Table 1** showcases a review of the deep learning infant brain segmentation literature along with limitations of deep learning algorithms. Each study reviewed has limited sample sizes of high-quality ground truth segmentations, with no study exceeding more than 25 infants. Most studies focus on limited age ranges, specifically between 5-9 months, limiting the generalizability of these models to other age groups. Outside of Moeskops et al ^33^, all studies within **Supplemental Table 1** also removed the skull, cerebellum, and brain stem from the images for training and prior to segmentation generation. This pre-processing likely reduces the generalizability for images in which the skull remains and increases the burden for performing inference. In addition, even in the case where sufficient data is used for training outputs often don’t follow common labels and standards such as the frequently used Freesurfer software. Last, implementing a model that is open-source, community-centered and follows Findable Accessible Interoperable and Reusable (FAIR) principles ^34^ is vital for the development and progress in future model generation and general usage.

BIBSNet (**B**aby and **I**nfant **B**rain **S**egmentation Neural **Net**work) provides a new open source software to advance brain segmentations. BIBSNet was implemented to add to and improve upon previous deep learning infant and baby brain segmentation research by creating a model that can handle variability of infant MR images based on age, neuronal developmental status, and data acquisition and quality. BIBSNet uses a larger sample size, n=90, than has been previously published and employs a relatively wide age range — 0-8 months. This wide age range allows BIBSNet to capably segment neonate and infant brains across important brain developmental boundaries (i.e., 0-3 months, 3-6 months, and 6-8 months). BIBSNet was built upon the proven deep learning architecture, nnU-Net ^35^, for brain segmentations. To improve generalizability and robustness, SynthSeg was used to create augmented images ^36^. Unlike models described in the previous literature, BIBSNet does not require that the skull, brainstem, or cerebellum be removed before segmentation; hence, much less pre-processing before inference is needed. The segmentations produced from BIBSNet are BIDS-compatible and conducted using input data in the subject’s native space. This choice was intentionally done so that BIBSNet segmentations can easily be transformed to fit whatever space is required for processing and analysis ^37^. BIBSnet is fully open-source, so other researchers are able to look at methodological detail and train their own models if needed. Finally, BIBSNet outputs can be directly input into a pre-processing pipeline, such as fMRIPrep-Infants ^38^ and the Developmental Cognition and Neuroimaging (DCAN) labs infant processing pipeline ^14,15,39,40^, in particular because it follows the standard Freesurfer anatomical labels, making it a turn-key solution for subsequent analyses.

## Results

### BIBSNet training is computationally expensive, but BIBSNet application is efficient

As one might expect, BIBSNet training necessitates considerable time and resources, requiring approximately 3.5 days of compute time with 1 NVIDIA V100 GPU, 6 CPUs, and 90GB of RAM. This resulted in a total clock time of 5-7 days factoring in average delays on the Minnesota Supercomputing Institute (MSI) systems. The training included 5 folds with 1000 epochs per fold, with each epoch taking around four to five minutes to complete. After an initialization step completed within the first fold, all five folds could be run concurrently.

While model training is resource-intensive, using the trained model for inference is straightforward. The BIBSNet application, provided as a BIDS app, requires minimal time and computing power to perform inference on unseen T1w/T2w image pairs. The inferences for the present study only required 4 minutes per subject using 2 CPUs and 20GB of RAM on the MSI systems. Details for downloading and usage can be found on Zenodo and GitHub^41^. This far out-performs the resources and time required by JLF, which, including the pre-processing steps required (i.e. nonlinear registration of a set of atlas images to the subject brain) can take up to 2-3 days.

### Dice similarity coefficient comparisons

DSCs were used to assess overlap in label assignments for BIBSNet- and JLF-derived segmentations compared to ground truth, i.e. manually corrected, segmentations for cortical and subcortical structures in the left and right hemispheres (**Figure 1**). For cortical structures, there was more similarity between BIBSNet and ground truth structures (RMSE = 0.157 gray matter, 0.144 white matter) than between JLF and ground truth (RMSE = 0.30 gray matter, 0.24 white matter), with an average DSC of 0.856 and 0.752 respectively across cortical structures. The DSCs were further compared with repeated sample T-test, revealing that both the gray matter and white matter DSCs produced from BIBSNet were significantly larger than JLF (gray matter: T = 14.08, p < 1×10^−23^; white matter: T = 6.21, p < 1×10^−7^). For subcortical structures, amygdala and hippocampus DSCs were compared as these are typically more difficult to accurately segment due to their small size. The bilateral amygdala and hippocampus RMSE based on the DSC for BIBSNet were RMSE = 0.17 and RMSE = 0.16, respectively, while for JLF they were RMSE = 0.24 and RMSE = 0.19. Neither subcortical comparison was statistically significantly (Amygdala: T = 1.74, p = 0.08; Hippocampus: T = 0.41, p = 0.67), however, there was much less variability in DSC values from BIBSNet vs ground truth (average standard deviation across regions = 0.04) compared to JLF vs ground truth (average standard deviation across regions = 0.12) (**Figure 1a**).

**Figure 1.**
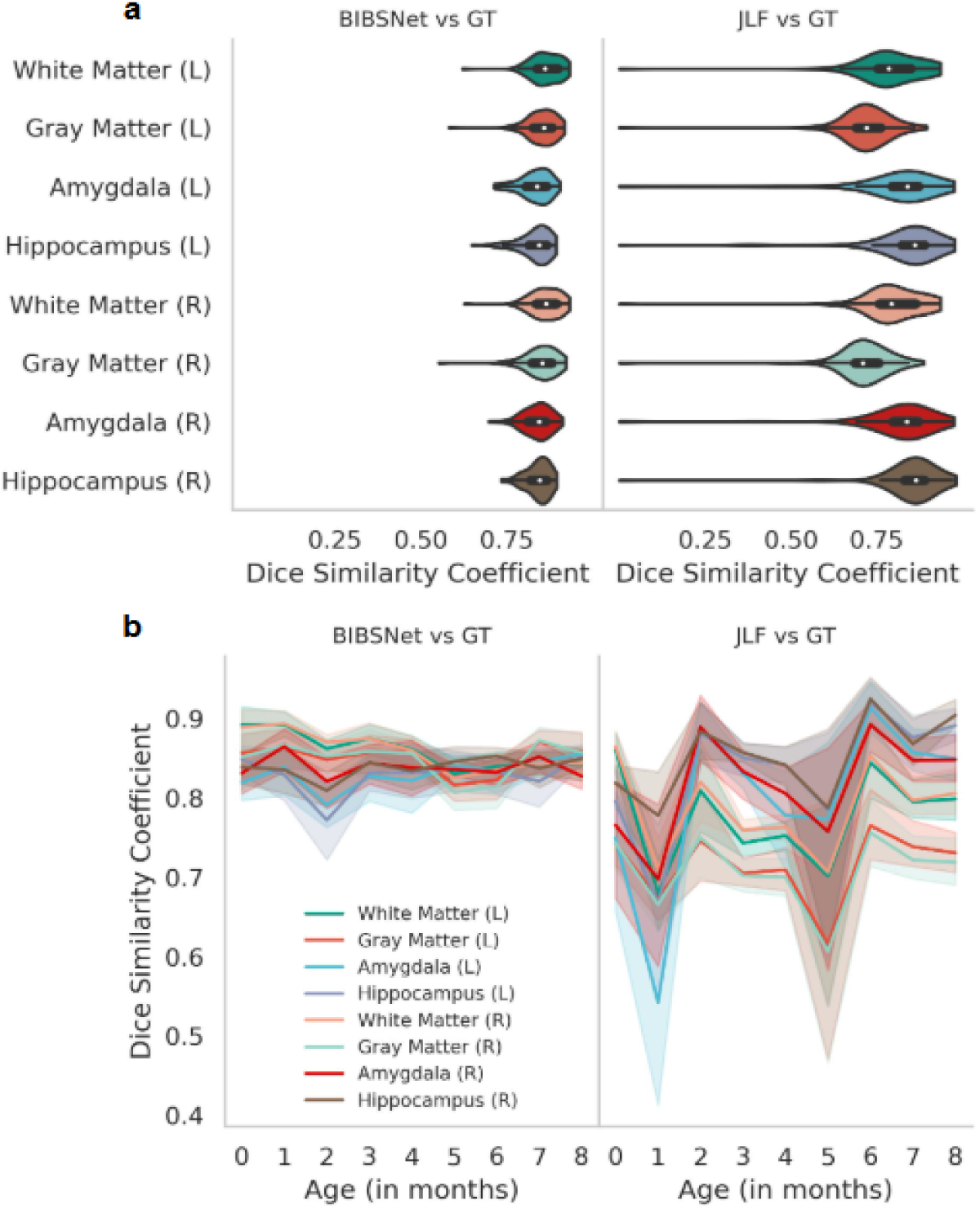
Dice similarity coefficients by infant age full sample, JLF and BIBSNet. **a**,**b** ALBERT and BCP analysis sample with inferred JLF and BIBSNet segmentations compared against Ground Truth (GT) using Dice Similarity Coefficient (DSC) on left and right white matter, left and right gray matter, left and right hippocampus, and left and right amygdala (n=90). **a,** Violin plots showcasing DSC between BIBSNet and GT segmentations (left), and JLF and GT segmentations (right). Notice that JLF performs worse compared to BIBSNet particularly the range of the data distribution. **b,** Line plots outlining the per subject DSC by infant age (in months) for BIBSNet (left), and JLF (right), and iBeat (right). The mean DSC is represented by opaque lines, whereas the semi-transparent lines show the 95% confidence interval variability. Observe, the large variability of DSC with JLF across infants of differing ages, whereas BIBSNet remains relatively stable across all ages.

An ad hoc comparison of BIBSNet to iBeat, an existing deep learning model for infant brain segmentation ^42,43^, was also performed. BIBSNet and iBeat differ, in that, BIBSNet produces FreeSurfer-compatible segmentations as well as segmenting cortical and subcortical structures, whereas iBeat does not produce FreeSurfer-compatible labels and segments the cortical structures only. Therefore, the analyses were limited to comparisons of white matter and gray matter. BIBSNet and iBeat performed similarly with no significant differences across ages within gray matter (T = 1.30, p = 0.19) (**Supplemental Figure 2a**). However, in white matter, while both performed similarly from 0-5 months (T = 0.69, p = 0.49), iBeat outperformed BIBSNet at months 6-8 significantly (T = 6.9, p < p < 1×10^−6^) (**Supplemental Figure 2b**).

### Volumetric, Cortical Surface, and Functional Connectivity Comparisons

We assessed the similarity of cortical and subcortical volumes in BIBSNet- and JLF-derived segmentations compared to ground truth segmentations. For both gray matter (BIBSNet RMSE = 38945.72, JLF RMSE = 42515.47) and white matter (BIBSNet RMSE = 38414.41, JLF RMSE = 47704.83), BIBSNet volumes were more similar to ground truth than JLF (**Figure 2a,c**). For subcortical volumes, we compared hippocampus and amygdala volumes to match the DSC comparisons above (**Figure 2b,d)**. Bilateral hippocampus volumes produced from BIBSNet (BIBSNet RMSE = 397.03) segmentations were more similar to ground truth than JLF (JLF RMSE = 465.72). Similarly, ground truth and BIBSNet bilateral amygdala volume were more similar (BIBSNet RMSE = 231.96) compared to JLF and ground truth (JLF RMSE = 237.51).

**Figure 2.**
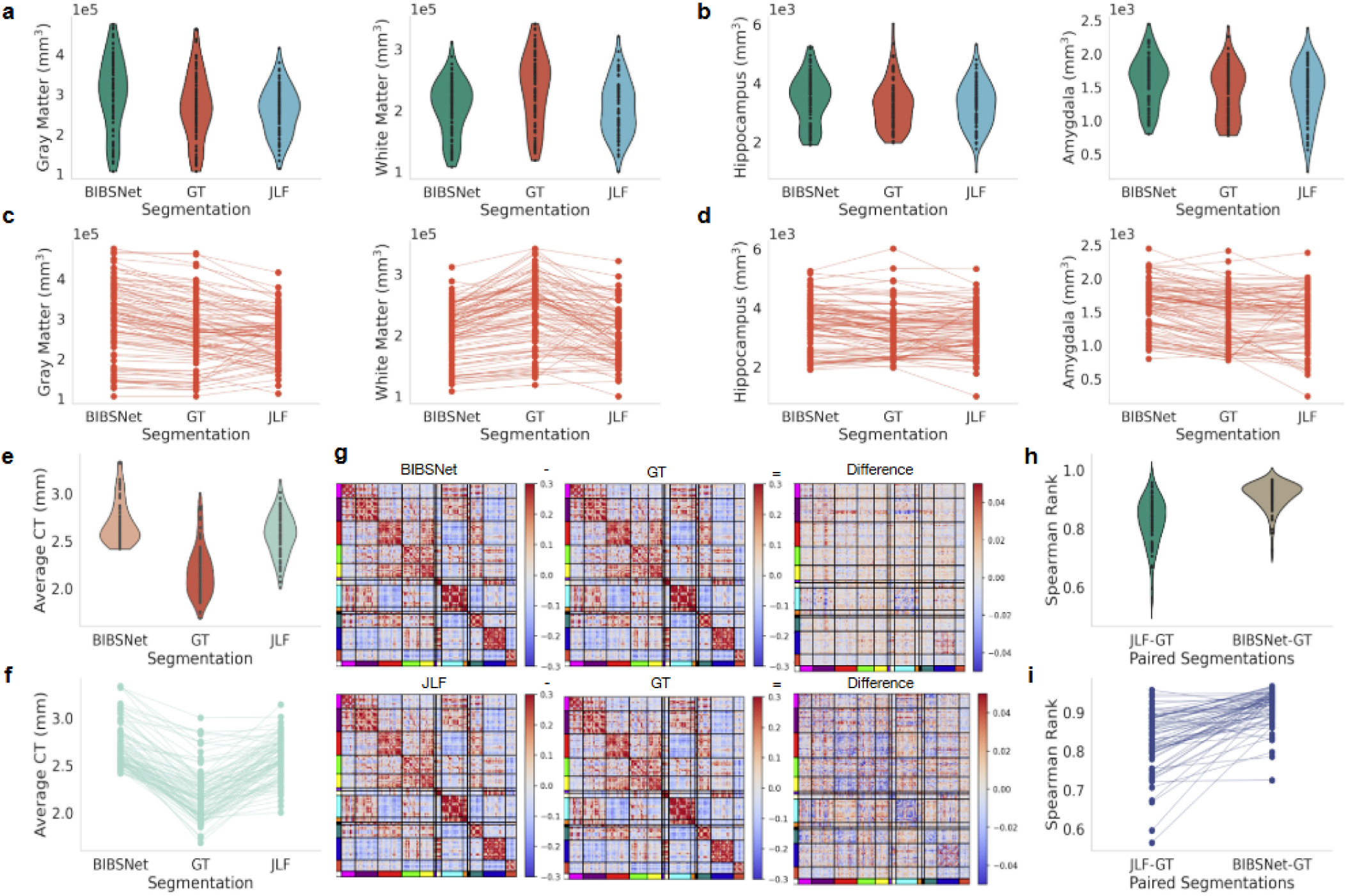
Anatomical and functional derived brain metric comparisons between JLF, BIBSNet and ground truth. BCP and ALBERT analysis sample, anatomical data: (a-d,f-h, n=90); functional data (e, n=64). a,c, Gray matter (left panels) and white matter (right panels) volume comparisons based on segmentations produced from BIBSNet, ground truth (GT), and JLF. b,d Bilateral hippocampus (left panels) and amygdala (right panels) volume comparisons based on segmentations produced from BIBSNet, GT, and JLF. a,b violin plots – a hybrid of a box and kernel density plot – showcasing the comparisons for each segmentation model grouping, c,d line plots showcasing the variability in cortical volume by segmentation model used, per participant. e,f, Cortical thickness values per surface vertex were calculated by FreeSurfer, averaged per surface, and plotted as violin plots (labeled as “Average CT”) (e) and line plots (f) showcasing the comparisons for each segmentation model grouping. g, Difference maps of segmentation group functional connectivity matrices derived from the Gordon parcellation atlas. Networks are marked in color along the X and Y-axes. BIBSNet (top) and JLF (bottom), separately were subtracted from by the Ground Truth functional connectivity matrix to produce a difference map. Notice that the values of the difference map between BIBSNet and GT hover closer to zero (i.e. no difference), compared to the JLF and GT difference map. h,i, Similarity from paired functional connectome matrices was calculated using Spearman’s rank correlation between JLF and GT and BIBSNet and GT and plotted as violin plots (h) and line plots (i). Notice that the distribution of values pile up near the correlation max (1.0) and is much tighter for BIBSNet compared to JLF.

Interestingly, although BIBSNet DSC and resulting cortical and subcortical volumes were more similar to ground truth, cortical thickness from JLF-derived segmentations were more similar to ground truth (RMSE = 0.45) than BIBSNet (RMSE = 0.55) (**Figure 2e,f**).

In addition to assessing the similarity between label assignments of segmentations and anatomical metrics derived from the associated MRI images, we also analyzed how functional connectivity compares between BIBSNet- and JLF-derived segmentations and ground truth. The similarity between BIBSNet and ground truth functional connectome matrices (RMSE = 0.09) was greater than the similarity between the JLF and ground truth (RMSE = 0.19) (**Figure 2g,h,i**).

## Discussion

BIBSNet is the only open-source pre-trained deep neural network model that we are aware of to construct high-quality cortical and subcortical segmentations in infant brains with 29 FreeSurfer compatible labels, highlighting its utility for use in infant processing methods and pipelines such as fMRIPrep-Infants^38^, MCRIBS^44^, and Infant FreeSurfer^45^. The extensive examination of BIBSNet showcases its success compared to JLF, the current state-of-the-art traditional approach, across DSC, morphometric, and functional metrics. An added benefit of BIBSNet is that it is nearly 600x faster than JLF and requires minimal computational resources. An ad-hoc examination comparing BIBSNet to iBeat, the current state-of-the-art deep neural network approach trained on infants, showed slightly higher DSC similarities. However, the comparison was limited to cortical structures as iBeat does not segment subcortical structures nor does it produce FreeSurfer compatible labels.

### BIBSnet segments cortical and subcortical ROIs with state-of-the-art accuracy

The analyses to evaluate the DSCs further confirms that BIBSNet segments cortical and subcortical structures with state-of-the-art accuracy. Prior literature DSC findings show that gray matter ranged from 0.84 - 0.92 with a mean of 0.879, while white matter ranged from 0.85 - 0.93 with a mean of 0.885^22,28,29,31,33,43,46–49^ (**Supplemental Table 1**), with BIBSNet firmly fitting into this. Across the cortical and subcortical ROIs evaluated with DSC BIBSNet outperformed JLF, the existing state-of-the-art traditional segmentation method (**Figure 1**). As showcased by the difference in standard deviation the distributions are much tighter for BIBSNet, highlighting its high reliability. Performance is also evident within the single subject comparisons as shown with **Figure 3**. Notice that there is high correspondence between the BIBSNet and ground truth segmentations more generally, whereas the arrows highlight the errors from the JLF segmentations. In addition to DSC comparisons with JLF, a comparison with iBeat, the existing state-of-the-art deep learning segmentation method trained on infants, was performed. iBeat and BIBSNet DSC comparisons revealed that the performance was similar between the two models, with iBeat slightly outperforming BIBSNet based on RMSE, and paired T-tests. It is worth highlighting that the differences between the means are quite small (gray matter = 0.007, white matter = 0.014), indicating that the differnences between BIBSNet and iBeat DSCs are subtle. When the DSC comparisons are observed at a per age level there is some performance variability between iBeat and BIBSNet. For example, iBeat outperforms BIBSnet in 2,3,5, and 6 month infants, whereas BIBSNet outperforms iBeat in 0-month and 7-month infants based on gray matter (**Supplemental Figure 1**). The similarity between BIBSNet and iBeat can also be seen within the single subject comparisons in **Supplemental Figure 2**. Importantly, the comparison of BIBSNet and iBeat is limited to cortical structures, as iBeat does not perform subcortical segmentations. BIBSNet has two important distinguishing features that stand out compared to the extant literature. First, nearly all models require pre-processing to remove part of the anatomy, particularly the skull. BIBSNet does not require image pre-processing, and is robust to the existence of the skull in images. Secondly, the training samples from models in prior literature involve infants between 5 and 9 months of age. BIBSNet, on the other hand, is trained on a much larger age range, between 0 and 8 months. These distinguishing characteristics allow BIBSNet to generalize well to unseen data.

**Figure 3.**
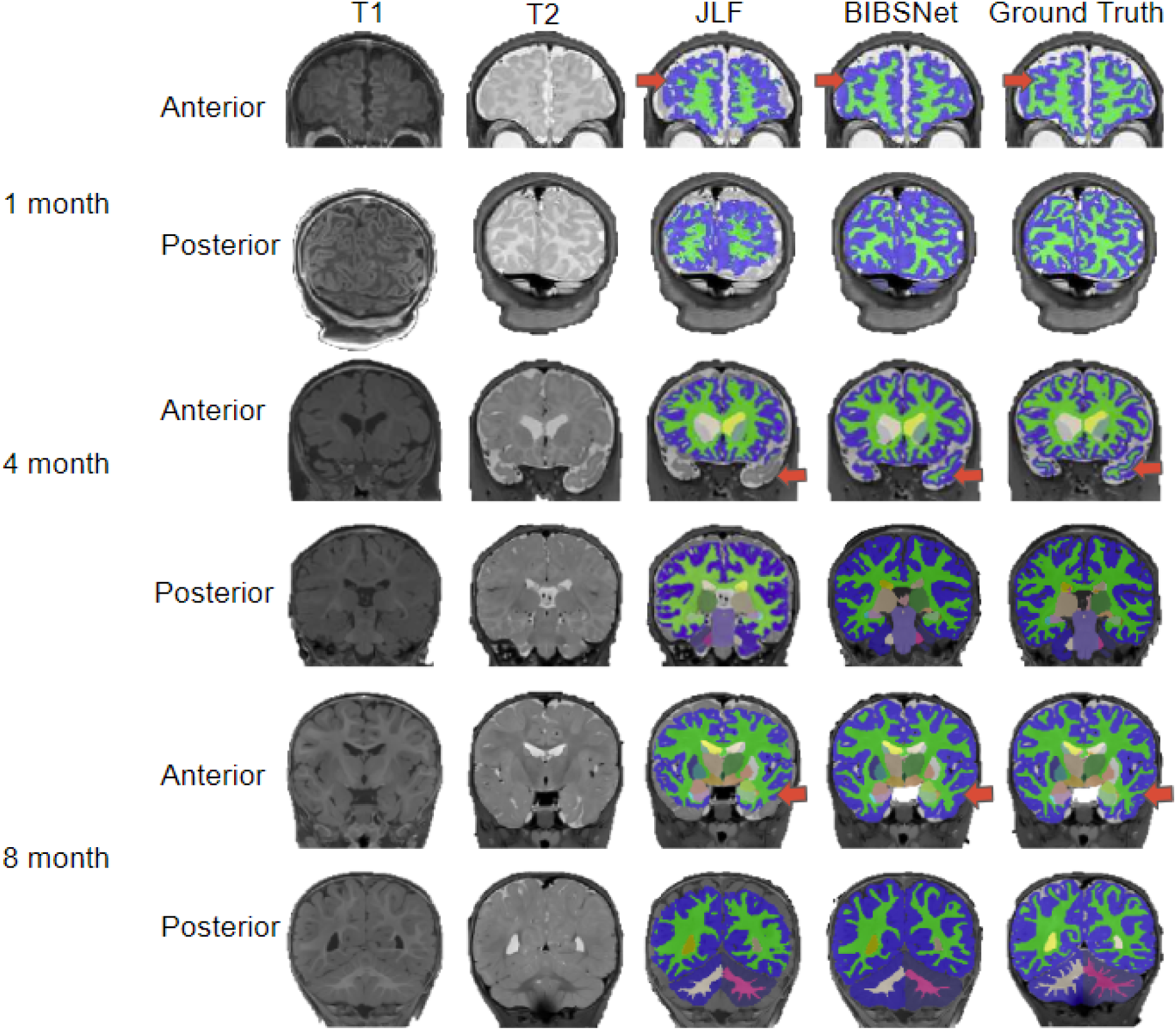
Comparison between BIBSNet, JLF, and ground truth segmentation with representative participants. Single subject representative slices on the anterior and posterior aspects for 1, 4, and 8 month infants showcasing the T1-weighted, and T2-weighted images, along with the segmentations produced from JLF, BIBSNet, and Ground Truth. The red arrows highlight segmentation label differences. In each case JLF fails to properly segment the white matter – lime green – or gray matter - dark blue.

### BIBSnet improves accuracy of brain morphometric and functional measures substantially over JLF

When the segmentations from ground truth, JLF, and BIBSNet were used to derive volumetric and cortical surface metrics, BIBSNet showed a similar pattern to the DSC comparisons. Across nearly all volumetric and cortical surface derived metrics, BIBSNet outperformed JLF. The effects were particularly pronounced with gray and white matter volume, with JLF showing nearly double the RMSE compared to BIBSNet (**Figure 2a,c**). Differences in RMSE highlights two details: 1) that JLF made more errors than BIBSNet; and 2) the errors that JLF made were markedly larger compared to BIBSNet. Both JLF and BIBSNet segment all subcortical structures, however, the amygdala and hippocampus were of particular focus as they are smaller structures and typically more difficult for automated methods to segment. Once again, BIBSNet outperformed JLF across both the hippocampus and amygdala. In both cases, it is likely that there is considerable variability across JLF template participants causing it to make some mistakes as it attempts to segment unseen data. On the other hand, BIBSNet seemed to have learned the latent patterns of each region allowing it to more accurately segment structures despite potential variabilities.

In addition to morphometrics being affected by the underlying brain segmentations, fMRI data produced from the DCAN infant pipeline is as well since the BOLD data is projected onto the cortical surface, a routine which requires a segmentation^14^. Therefore, functional connectivity matrices were compared across segmentation type. Using Spearman’s rank correlation to generate a measure of similarity of paired functional connectivity matrices, RMSE revealed that functional connectome matrices derived from BIBSNet segmentations were much more similar to ground truth than JLF derived matrices (**Figure 2 h,i**). This effect is also apparent within the group averaged based difference matrices (**Figure 2g**), notice that values from the BIBSNet and ground truth difference matrix hovering closer to zero, indicating no difference, compared to the JLF and ground truth difference matrix.

### Impact and Future directions

BIBSNet provides a critical need for infant neuroimaging studies. Many neuroimaging pipelines ^44,45,50,51^ require FreeSurfer compatible labels from segmentations, a feature which is unavailable with existing deep learning based infant segmentation methods. With the use of FreeSurfer compatible labels in BIBSNet, it is an easy turnkey solution for incorporation into existing pipelines and can be relied upon for segmentation within yet to be designed pipelines. The addition of subcortical segmentations in BIBSNet make it invaluable for handling multimodal infant studies, such as HBCD. For example, subcortical segmentation is needed for diffusion^51^ and MRS^52^ data processing. Moreover, it is critical for deriving measures from fMRI for subcortical structures^50^ as well. Additionally, our hope is that the morphological and resting state fMRI analysis performed, paired with the images showcased within **Figure 3** indicating the per subject improvement in accuracy of BIBSNet over JLF, will allow researchers to see the practical benefits of using BIBSNet. Finally, as a resource the BIBSNet lookup table is provided in **Supplemental Table 2** so others can easily know the mapping between label numbers within the segmentations and their corresponding label names.

Given the success of BIBSNet, future directions are warranted. A primary feature of the BIBSNet model proposed in this work is that it currently requires a T1w and T2w image pair. In practice, infant MRI research studies typically only acquire one of the two anatomical images. Since infant MR scans are typically acquired while the infant is asleep, time is at a premium.

Therefore, once a single anatomical image is collected, researchers often move on to collecting other images such as resting state or diffusion weighted imaging. As a result of this, we have trained additional BIBSNet models, one that requires just a T1w and the other just a T2w image. A future study will evaluate the performance of these additional BIBSNet models. Additional, future work will involve the incorporation of large and diverse datasets, such as HBCD, to widen the age range and acquisition parameters of images in the training sample. This will boost model generalizability for out-of-sample inference and increase its utility for researchers that collect data with a variety of ages and/or acquisition parameters.

## Conclusion

In conclusion, BIBSnet shows state-of-the-art performance when compared to iBeat, JLF and models used in prior literature. BIBSNet performed well across multiple ages and was robust to the existence of the skull in images. Finally, the pre-trained model is fast (at least 600x speed up compared to JLF), requires minimal high performance computing resources, and can easily be included in other pipelines. Future studies will extend BIBSnet’s performance and flexibility across more datasets and longitudinal timepoints.

## Methods

**We used MR images and manually annotated segmentations from infants aged 0 to 8 months from the BCP and ALBERTs datasets for the present study.**

### MR Image Collection Procedure

MR images were collected from 72 participants within the BCP study^53^ — median age at scan = 5.5 months, 43 female — with a 3T Siemens Prisma at the University of Minnesota’s Center for Magnetic Resonance Research (CMRR). Images used for this study included T1-weighted (echo time = 2.24 ms, repetition time = 2400 ms, inversion time = 1060 ms, sagittal slices = 208, flip angle = 8°, matrix = 320 x 320, voxel sizes = 0.8 x 0.8 x 0.8mm^3^), T2-weighted (echo time = 564 ms, repetition time = 3200 ms, sagittal slices = 208, flip angle = variable, matrix = 320 x 320, voxel sizes = 0.8 x 0.8 x 0.8mm^3^), and approximately 12 minutes of resting state MR images (echo time = 37 ms, repetition time = 800 ms, axial slices = 72, flip angle = 52, matrix = 104 x 91, multi-band acceleration factor = 8, voxel sizes = 2.0 x 2.0 x 2.0mm^3^) made up of two separate collections with reverse phase encoding (AP and PA).

To supplement the sample with additional neonates, 19 infants that were used to generate the ALBERTs neonatal atlas were also used ^54^. The MRI data for the ALBERTs infants (median age at scan = 0 months) were acquired on a 3.0 T Philips Intera scanner (Philips Medical Systems, Best, Netherlands). The relevant images utilized included T1-weighted (echo time = 4.6ms, repetition time = 17 ms, sagittal slices = 124–150, flip angle = 30°, matrix = 256 × 256, voxel sizes = 0.82 × 0.82 × 1.6 mm^3^) and T2-weighted (echo time = 160ms, repetition time = 8000 ms, sagittal slices = 88–100, flip angle = 90°, matrix = 224 × 224, voxel sizes = 0.86 × 0.86 × 2.0 mm^3^) data.

### Manually Annotated Segmentation Procedure

Following data collection, the T1w and T2w MR images from the BCP study were used to produce manual annotations based on FreeSurfer’s aseg atlas^55^. As a starting point for the segmentations, the T1w and T2w images from each BCP participant were run through either JLF or a prototyped version of the BIBSnet algorithm. Led by an expert (E.F.), the cortical and subcortical structures from the BCP participants were then extensively manually annotated with highly trained staff to generate accurate segmentations. This work, known as the Baby Open Brains (BOBs) repository, will be made available to others as a resource^56^. The ALBERTs MR images were included in the training to have fairly equal sample sizes of infants from 0 to 8-months. The segmentations produced from the ALBERTs MR images had been previously manually annotated but did not have extensive quality control procedures and were not reviewed by an expert.

The BCP and ALBERTs MR images and manual annotated segmentations were used for BIBSNet model training and subsequent data processing and analyses.

**BIBSNet combines the nnU-Net model and SynthSeg software, to produce generalizable segmentations**

### nnU-Net model design

As can be seen within **Figure 4**, nnU-Net plays a central role in the BIBSNet model design. nnU-Net is a model that is based upon the CNN based network, U-Net ^57^. In the last several years, U-Net has shown state-of-the-art performance in segmentation problems, specifically biomedical image segmentation. U-Net consists of two separate elements, the contracting path and the expanding path. The contracting path progressively downsamples the input image/s while gaining progressively more in-depth features, known as feature maps, to represent the image. Following the contracting path, the expansive path takes the feature maps and progressively up-samples them until they are the same size as the originally input image/s. The downsampling and upsampling is what gives this model its characteristic “U” shape, hence the “U” in U-Net. Despite the success of U-Net, a major pitfall was its difficulty with applying it to different image analysis problems. It required a multitude of expert decisions in the form of adaptation and modification of parameters to be made without clear guidance or obvious parameter defaults. nnU-Net sought to correct this by 1) fixing parameters that do not require adaptation; 2) tuning parameters that needed to be adapted based upon the inputted dataset; and 3) for any parameters that remain, making decisions empirically from the data ^35^.

**Figure 4.**
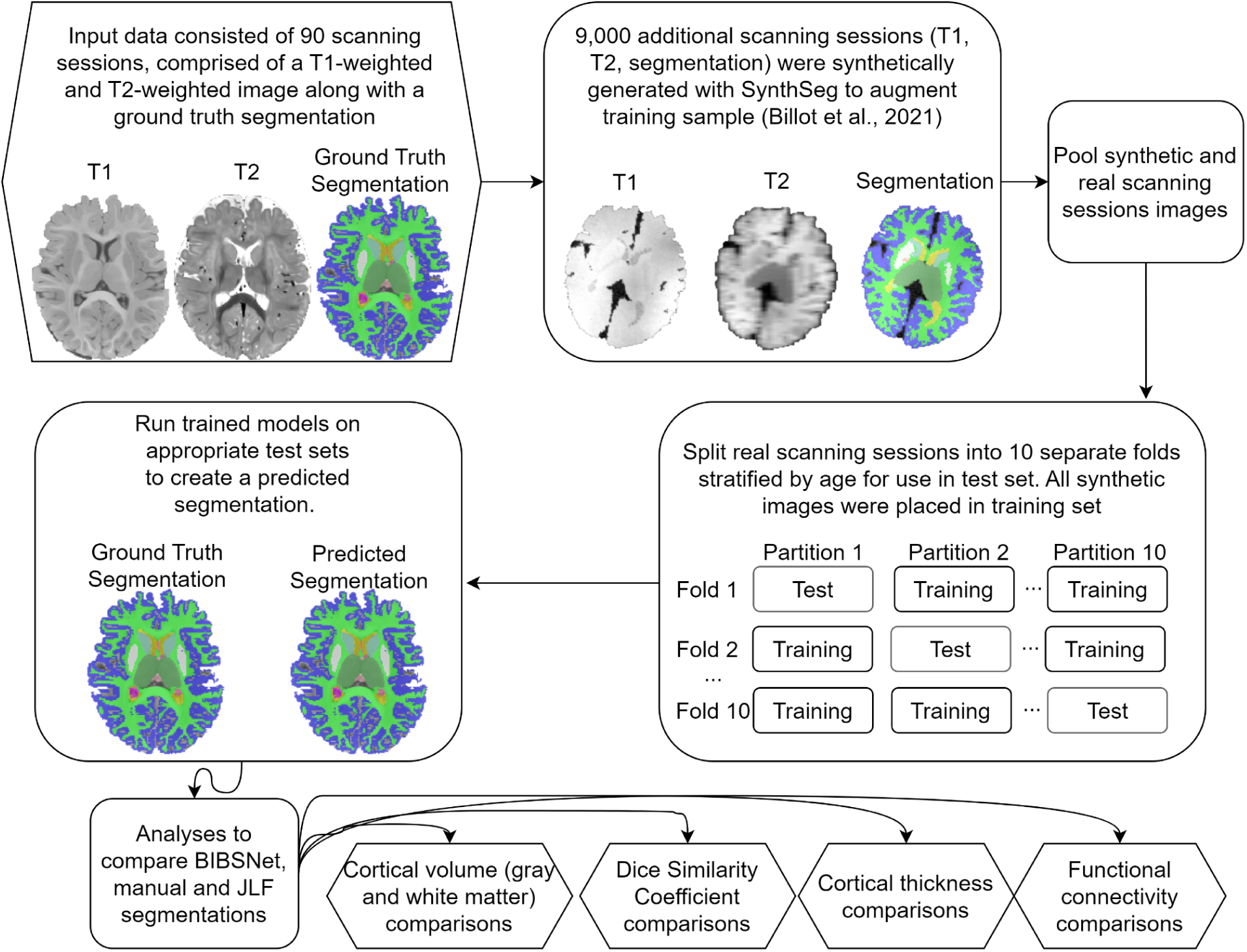
Schematic of model training, cross validation, and evaluation process. Schematic overview of BIBSNet model design, training, and cross validation process. Sections shaped as hexagons represent the starting and end points of the current work, whereas sections shaped as rounded rectangles represent intermediate steps.

### SynthSeg

While nnU-Net plays the integral role of segmenting the inputted images, SynthSeg is vital to improve the generalizability of the model. SynthSeg is software that is capable of producing synthetic MR imaging data. Here, the manually annotated segmentations from the BCP and ALBERTs were inputted into SynthSeg for the production of synthetic MR images. SynthSeg randomly modified the features of each image in four separate ways to emulate different acquisition parameters and data quality ^36^. First, an affine and non-linear transformation was applied to spatially deform the data, simulating a head tilt or changes in head size. Second and third, a randomized Gaussian mixed model and bias corruption were applied. These steps are meant to mimic the global and local pixel intensity differences that can occur across different MRI scanners, such as bias field artifacts. Fourth, to mirror acquisition differences resulting from slice thickness, collection orientation, or voxel size the image was randomly downsampled and blurred. For the current study, 9,000 T1w, T2w, and segmentation datasets were produced based on the 90 participant datasets, a 100x increase in training data. Henceforth, images from the BCP and ALBERTs will be referred to as “real images”, while the synthetic images produced by Synthseg will be referred to as “synthetic images”.

### BIBSNet cross validation and training

As shown in **Figure 4**, the synthetic outputs from SynthSeg, in combination with the real BCP and ALBERTs MR imaging data, were fed directly into nnU-Net for segmentation. Importantly the imaging data were not skull-stripped, as a desired feature was for BIBSNet to perform inference on non-skull stripped images. The same BCP and ALBERT participants were used for training and analysis, thus, a k-fold training strategy was employed to avoid data leakage. The first step included splitting the real images into ten approximately equally sized pseudorandom testing datasets stratified by age. Each of the ten testing datasets was paired with a training dataset. The training dataset was composed of the other real images along with all generated synthetic images. This strategy ensured that when inference was performed the real images were previously unseen by at least one fold.

To avoid chirality errors, where the inferred annotation was locally correct but the hemisphere was not (e.g., right white matter labeled on the left hemisphere), inferred nnUnet segmentation outputs underwent a chirality correction procedure. The anatomical T1 was non-linearly registered via ANTS SyN algorithm^58^ to a template infant T1 of the same age. The template infant comprised a mask of the left and right hemisphere mapped to the T1. The non-linear warp was inversely applied to the left/right mask using a nearest neighbor interpolation, producing the left/right mask in the BIBSnet segmentation space. The mask was then applied to the BIBSnet segmentation, correcting any chirality errors.

**MRI data were processed with the DCAN lab’s infant-ABCD-BIDS processing pipeline using segmentations produced from ground truth and the JLF and BIBSNet models**

To test for the effect of segmentation type in subsequent analyses, each of the 90 participants were processed through the DCAN labs infant processing pipeline^14,39,40,59^ in three separate ways: 1) running JLF using the age-matched templates produced from the manually annotated segmentations (excluding the same-subject template) 2) using the externally generated segmentations based on the manual annotations; and 3) using the externally generated segmentation inferred by the trained BIBSNet model. From here on, the processing strategies will be referred to as “JLF”, “ground truth” and “BIBSNet”, respectively. Beyond the three separate processing strategies mentioned above, the data were processed using the exact same procedures. iBeat inferred segmentations were not included in this processing, in part, because iBeat does not segment subcortical structures so there was concern regarding the ability to process the data and the accuracy of the potential outputs. However, iBeat was included in Dice Similarity Coefficient (DSC) comparisons.

The DCAN lab’s infant processing pipeline is based on the Human Connectome Project minimal processing pipeline^14^ with key changes to accomodate for the differences in the developing brain of infants. Additionally, the design of DCAN lab’s infant processing pipeline draws heavily on ABCD-BIDS processing pipeline to accomodate and attenuate scanner effects found within major MRI vendors GE, Philips, and Siemens ^16^. The pipeline is split into several stages: “PreFreeSurfer”, “FreeSurfer”, “PostFreeSurfer”, “FMRIVolume”, “FMRISurface”, “DCANBOLDProcessing”, and “ExecutiveSummary”.

The “PreFreeSurfer” stage, the first stage in the structural pipeline, aligned the T1-weighted and T2-weighted anatomical images, registered native structural images to standard MNI space, and, in the case of JLF processing, ran JLF. The second stage, “FreeSurfer”, ran the software tool FreeSurfer ^13,55,60–64^. The primary goals of FreeSurfer are to reconstruct white and pial cortical surfaces, segment subcortical structures, and to register produced native surfaces to the template space, fsaverage. “PostFreeSurfer”, generated the NIFTI volumes and GIFTI surface files and placed them together into CIFTI files for viewing in the visualization tool Connectome Workbench ^14^.

The main goals of the “FMRIVolume’’ stage, the first in the functional pipeline, are to remove spatial distortion, realign volumes to correct for participant motion, register the functional to the structural data, reduce bias field, normalize the resulting 4D image to a global mean, and mask the data ^14,16^. The second functional pipeline stage is “FMRISurface”. The purpose of “FMRISurface’’ is to extract the time series processed within the volume space and place it into the surface CIFTI standard space. Voxels on the cortical gray matter ribbon were mapped onto the native cortical surface, transformed according to the surface registration onto the 32k mesh. Subcortical gray matter voxels were mapped from the set of pre-defined subcortical parcels to a standard set of voxels in each atlas parcel ^14,16^. For infant data, the standard adult MNI template cannot be non-linearly registered for subcortical data. Therefore, a piecemeal approach was used, where we register each subcortical region linearly to the adult template and then project the subcortical fMRI data into the CIFTI subcortical greyordinates.

Immediately following “FMRISurface”, two additional stages “DCANBOLDProcessing” and “ExecutiveSummary” were launched. The primary goals of “DCANBOLDProcessing” are to further process fMRI CIFTI data by filtering motion estimates to separate true head motion from factitious motion due to magnetic field changes due to breathing^15^ and produce both dense (dtseries) and parcellated (ptseries) CIFTI files for subsequent analyses. The final stage in the DCAN labs infant processing pipeline, “ExecutiveSummary”, summarized standard quality control outputs and provided them in a browser-interface for easy navigation and review ^16^. Data processing failed for a total of three participants. Two participants were from ALBERTs and one was from the BCP study. All three failures used the JLF inferred segmentations. These failures were removed from subsequent analyses, including cortical thickness comparisons and, in the case of the BCP, participant functional connectome and DSC analyses. Further details on failures are highlighted in **Supplemental Table 3**.

**Segmentation overlap and structural and functional MRI metrics were used to perform statistical analyses comparing JLF versus ground truth, iBeat versus ground truth and separately, BIBSNet versus ground truth**

#### Segmentation similarity analysis

To evaluate how similar the segmentations produced by JLF and BIBSNet were to the ground truth segmentations, DSC was used ^65,66^. DSC measures the fraction of overlap between anatomical regions of the same type, ranging in values between 0 and 1. DSC was calculated thrice, first to compare JLF and ground truth, second comparing BIBSNet and ground truth, and third, as a post-hoc analysis, to compare iBeat and ground truth. As iBeat does not segment subcortical structures, manually annotated images were used to mask out the would-be subcortical structures.

To assess DSC for each segmentation type, Root Mean Squared Error (RMSE) to the ground truth was used. RMSE is a metric often used to assess the accuracy of predictive models by ascertaining how closely predicted values match actual values. It is defined as the square root of the average squared differences between predicted and actual values. The lower the value of the RMSE, the better performance of the model. For the comparison, since DSC is a pairing metric, the ground truth DSC was a perfect 1.0 as each segmentation was compared with itself. RMSE, compared to other metrics like mean absolute error, punishes larger errors. We find that this feature is particularly useful for our comparisons as even subtle differences in derived metrics could result in different conclusions when studied or analyzed, let alone patently large errors. As an added assessment of method effectiveness paired T-tests are also used.

#### Structural MRI analyses

In addition to evaluating the similarity of resulting segmentations, morphological metrics were also analyzed. Using outputs from the FreeSurfer processing stage, four different morphological metrics – gray matter, white matter, bilateral amygdala, and bilateral hippocampal volume – and one cortical sheet metric – average cortical thickness – were analyzed. To assess method effectiveness, the metrics produced from BIBSNet segmentations were directly compared RMSE to the ground truth segmentation produced metrics. Separately, the metrics produced from the JLF model were directly compared to ground truth with the same strategy.

#### Functional MRI analysis

In addition to structural or structurally derived metrics, functional MRI metrics were also analyzed. The Gordon^67^ cortical parcellated resting state time series generated by the DCANBOLDProcessing processing stage were used to produce pair-pair correlation matrices with the tool “cifti-connectivity” (https://github.com/DCAN-Labs/cifti-connectivity). First, “cifti-connectivity”, used the Gordon cortical parcellated resting state time series processed through the DCAN labs infant processing pipeline. Second, timepoints were regressed out that exceeded a framewise displacement of 0.3mm ^68^. Regressing out time points in this fashion had the effect of removing sufficiently large motion events that could cause spurious correlation.

With these motion events removed, each so-called “filtered” Gordon parcellated resting state time series was temporally correlated with itself to produce a pair-pair correlation matrix, also known as a functional connectome matrix.

Like all other metrics, the functional connectome matrices were produced based on each of the three processing strategies. Spearman’s rank correlation was calculated per participant between the ground truth and BIBSNet and, separately, ground truth and JLF. This created a measure of per participant similarity between the JLF and ground truth, and BIBSNet and ground truth functional connectome matrices. Spearman’s rank correlation ranges from 0.0, meaning absolutely no similarity, and 1.0 meaning perfect rank similarity. RMSE was also calculated for comparisons. Similar to DSC, Spearman’s rank is a paired metric, thus the ground truth similarity had a perfect 1.0 correlation as each ground truth functional connectome was compared to itself.

## Supplementary Information

**Supplemental Table 1.** Spreadsheet of previous literature. Sample size, deep learning architectures, anatomy removed and Dice Similarity Coefficient reported in the recent infant brain segmentation literature. GM: gray matter, WM: white matter, CB: cerebellum, mWM: myelinated white matter, BGT: basal ganglia and thalami, vCSF: ventricular cerebrospinal fluid, uWM: unmyelinated white matter, BS: brain stem, cGM: cortical gray matter, eCSF extracerebral cerebrospinal fluid

| Authors , Year | Paper Title | Test Sample Size | Deep Learning Architecture | Anatomy Removed from Image | Dice Similarity Coefficient |
| --- | --- | --- | --- | --- | --- |
| Zhang, 2015 | Deep convolutional neural networks for multi-modality isointense infant brain image segmentation | 8: 26-34 wks | Convolutional Neural Network | skull, cerebellum, brain stem | GM: 0.86, WM: 0.85 |
| Moeskops, 2017 | Isointense infant brain MRI segmentation with a dilated convolutional neural network | 23: 26 wks | Convolutional Neural Network | skull, cerebellum, brain stem | WM: 0.874, GM: 0.877 |
| Moeskops, 2016 | Automatic Segmentation of MR Brain Images with a Convolutional Neural Network | 10: 30 wks<br>12: 40 wks | Convolutional Neural Network | None | 30 weeks: CB: 0.92, mWM: 0.69, BGT: 0.92, uWM: 0.96, BS 0.87, cGM: 0.84,<br>40 weeks: CB: 0.93, mWM: 0.56, BGT: 0.91, uWM: 0.93, BS: 0.85, cGM: 0.87 |
| Nie, 2016 | FULLY CONVOLUTIONAL NETWORKS FOR MULTI-MODALITY ISOINTENSE INFANT BRAIN IMAGE SEGMENTATION | 10: 26-34 wks | multi fusion fully convoluted network | skull, cerebellum, brain stem | GM: 0.873, WM: 0.887 |
| Dolz, 2018 | ISOINTENSE INFANT BRAIN SEGMENTATION WITH A HYPER-DENSE CONNECTED CONVOLUTIONAL NEURAL NETWORK | 23: 26 wks. | 3D hyper dense CNN known as HyperDenseNet | skull, cerebellum, brain stem | GM: 0.920, WM: 0.901 |
| Dolz, 2020 | Deep CNN ensembles and suggestive annotations for infant brain MRI segmentation | 23: 26 wks | 3D CNN SemiDenseNet, extending HyperDenseNet. | skull, cerebellum, brain stem | GM: 0.92, WM: 0.90 |
| Nie, 2019 | 3-D Fully Convolutional Networks for Multimodal Isointense Infant Brain Image Segmentation | 11: 26-34 wks | 3D multimodal fully convoluted network | skull, cerebellum, brain stem | GM: 0.8817, WM: 0.8586 |
| Zeng, 2018 | MULTI-STREAM 3D FCN WITH MULTI-SCALE DEEP SUPERVISION FOR MULTI-MODALITY | 23: 26 wks | 3D fully convoluted network | skull, cerebellum, brain stem | GM: 0.916, WM: 0.896 |
|  | ISOINTENSE INFANT<br>BRAIN MR IMAGE<br>SEGMENTATION |  |  |  |  |
| Wang,<br>2018 | Volume-Based Analysis of<br>6-Month-Old Infant Brain<br>MRI for Autism Biomarker<br>Identification and Early<br>Diagnosis | 18: 26 wks | Anatomy-Guided<br>Densely-Connected U-Net | skull | GM: 0.923, WM: 0.933 |
| Wang,<br>2023 | iBEAT V2.0: a<br>multisite-applicable, deep<br>learning-based pipeline for<br>infant cerebral cortical<br>surface reconstruction | 505: 29-45<br>postmenstrual<br>weeks | Anatomy-Guided<br>Densely-Connected U-Net | Skull, cerebellum, brain<br>stem | GM: ~0.86, WM: ~0.90 |

**Supplemental Figure 1.**
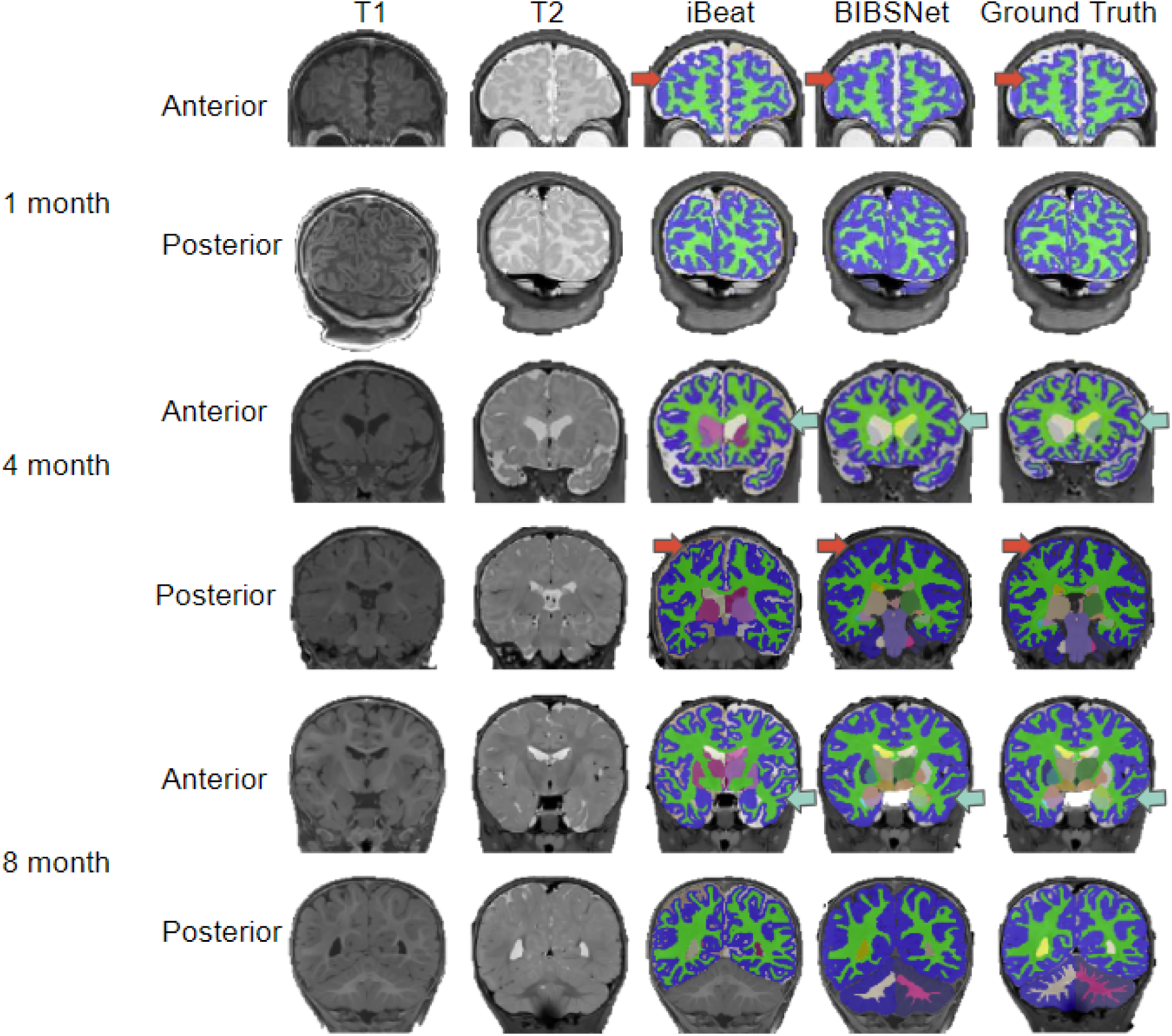
Comparison between BIBSNet, iBeat, and ground truth segmentation with representative participants. Single subject representative slices on the anterior and posterior aspects for 0, 4, and 8 month infants showcasing the T1-weighted, and T2-weighted images, along with the segmentations produced from iBeat, BIBSNet, and ground truth. The red arrows and light green arrows highlight segmentation label differences, whereby red arrows indicate where iBeat does better, and light green arrows where BIBSNet does better.

**Supplemental Figure 2.**
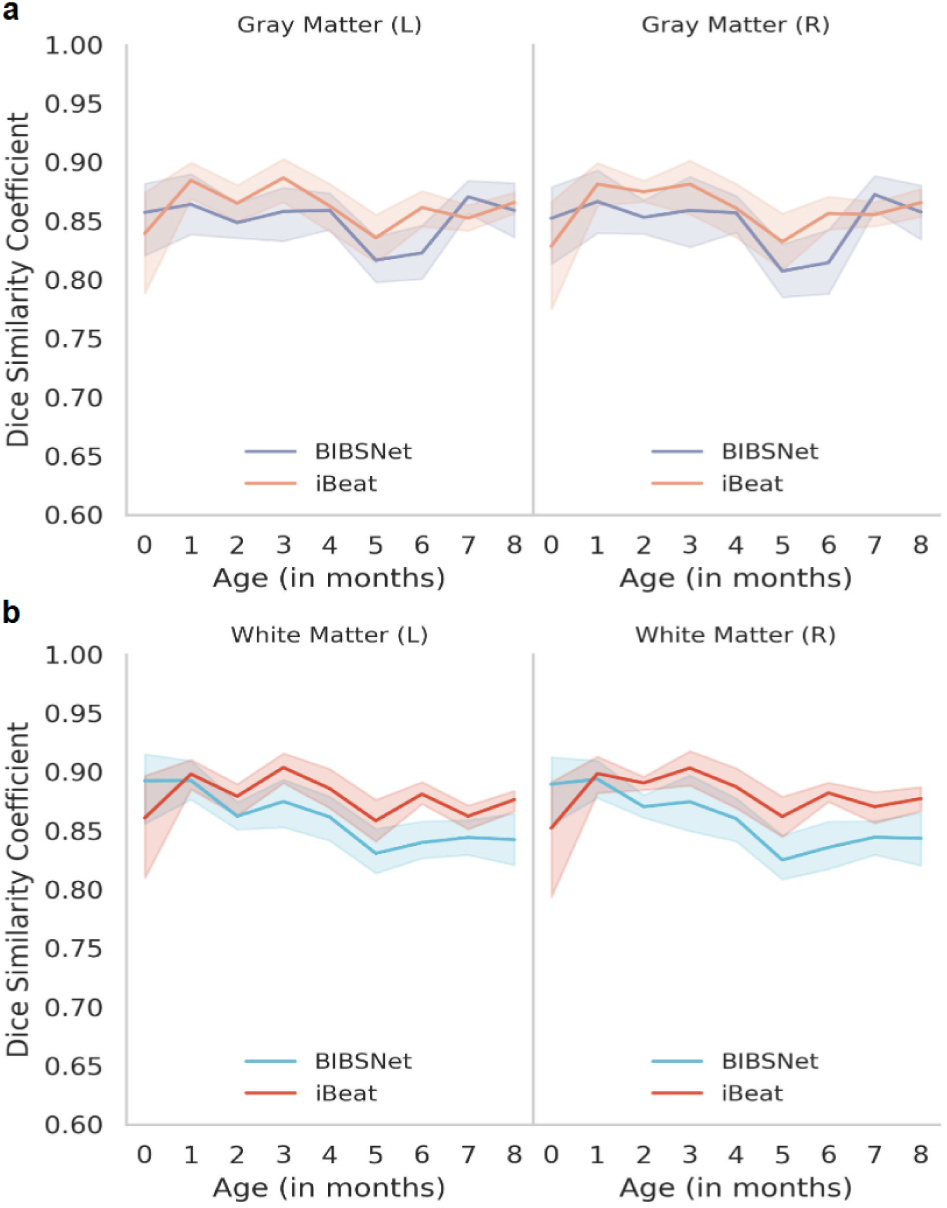
Dice similarity coefficients by infant age full sample, BIBSNet and iBeat. **a,b** ALBERT and BCP analysis sample with inferred iBeat, and BIBSNet segmentations compared against ground truth using Dice Similarity Coefficient (DSC) on left and right gray matter (**a**) and white matter (**b**) (n=90). Line plots outlining the per subject DSC by infant age (in months) for BIBSNet and iBeat. The mean DSC is represented by opaque lines, whereas the semi-transparent lines show the 95% confidence interval variability. Notice that BIBSNet and iBeat remain relatively stable across all ages. The DSC values diverge most heavily in 0-month infants (BIBSNet better), and 5-month infants (iBeat better). **a**) iBeat DSC = 0.856, iBeat RMSE = 0.151, BIBSNet DSC = 0.849, BIBSNet RMSE = 0.157, **b**) iBeat RMSE = 0.134, iBeat DSC = 0.876, BIBSNet RMSE = 0.144, BIBSNet DSC = 0.862

**Supplemental Table 2.** BIBSNet Lookup Table. Lookup table for the labels that BIBSNet segments. The label number represents the value that will be found within the segmentation images, whereas, the label name is the actual structure/region that the label number corresponds to.

| Label Number | Label Name |
| --- | --- |
| 2 | Left Cerebral White Matter |
| 3 | Left Cerebral Cortex |
| 4 | Left Lateral Ventricle |
| 8 | Left Cerebellum Cortex |
| 10 | Left Thalamus Proper |
| 11 | Left Caudate |
| 12 | Left Putamen |
| 13 | Left Pallidum |
| 14 | 3rd Ventricle |
| 15 | 4th Ventricle |
| 16 | Brain Stem |
| 17 | Left Hippocampus |
| 18 | Left Amygdala |
| 24 | CSF |
| 26 | Left Accumbens Area |
| 28 | Left Ventral Diencephalon |
| 41 | Right Cerebral White Matter |
| 42 | Right Cerebral Cortex |
| 43 | Right Lateral Ventricle |
| 47 | Right Cerebellum Cortex |
| 49 | Right Thalamus Proper |
| 50 | Right Caudate |
| 51 | Right Putamen |
| 52 | Right Pallidum |
| 53 | Right Hippocampus |
| 54 | Right Amygdala |
| 58 | Right Accumbens Area |
| 60 | Right Ventral Diencephalon |
| 172 | Vermis |

**Supplemental Table 3.** Processing Pipeline Failures. Table laying out failures through the DCAN infant pipeline. The left column highlights the segmentation type that failed. Middle column indicates what stage of the DCAN infant pipeline failed. Right column indicates the reason for the failure.

| Segmentation Type | Failure Stage | Reason |
| --- | --- | --- |
| JLF | FreeSurfer | Cortical Reconstruction Step Failed |
| JLF | FreeSurfer | Cortical Reconstruction Step Failed |
| JLF | PreFreeSurfer | JLF was unable to generate a segmentation |

## Acknowledgements

- The authors acknowledge the Minnesota Supercomputing Institute (MSI) at the University of Minnesota for providing resources that contributed to the research results reported within this paper. URL: http://www.msi.umn.edu
- The authors acknowledge the Center for Magnetic Resonance Research (CMRR) at the University of Minnesota for providing resources for MRI data collection. The MRI data collection was supported by grants NIBIB P41 EB027061 and 1S10OD017974-01 “High Performance Connectome Upgrade for Human 3T MR Scanner”
- This work was supported by National Institutes of Health grants U01 MH110274, R01 MH104324, P50 HD103573, U01 DA055371, U01DA041148, DA050291, MH096773 MH125829, DA055330 and the Bill & Melinda Gates Foundation INV-015711, author JM was funded by the DFG German Research Foundation 493345456 (Deutsche Forschungsgemeinschaft)

